# Parallel formation of opposing memories tunes online and pre-emptive control of learned behavior in eyeblink conditioning

**DOI:** 10.1101/2025.05.26.656060

**Authors:** Ryota Iwase, Shin-ya Kawaguchi

**Affiliations:** Department of Biophysics, Graduate School of Science, Kyoto University; Kyoto, 606-8502, Japan

## Abstract

Animals change their behavior and adapt to the environments through associating patterns of sensory experiences, making memories to acquire and/or cease actions. However, it remains to be clarified how distinct memories are formed and interact in response to altered sensory inputs. Here we studied this issue using contrasting paradigms for eyeblink conditioning in mice, and demonstrated a fundamental mechanism of memory counteraction to fine-tune the adaptive change of animal behavior. Excessive neutral sensory stimulation intermingled with the paired presentation of two sensory stimuli did not disturb the acquisition of conditioned responses, but gradually suppressed the occurrence of learned behavior during individual daily sessions. Modeling of acquisition and extinction processes, coupled with experimental validation of a model prediction, unveiled independent and parallel formation of long-term opposing memories for execution of eyeblink and its suppression irrespective of the temporal order: the memory to suppress the conditioned response can, surprisingly, be formed even before its acquisition. Consequently, the negative memory formed either online or pre-emptively calibrates the acquired behavior, optimizing motor performance reflecting both positive and negative sensory histories.

## Introduction

Animals adapt to changing environments by learning from experience, whose richness has been thought to affect the efficiency of adaptive behavior (Grant & Shipper, 1952; Thomas & Wagner, 1964). Notably, animals need to adaptively change the behavior even through a survival-related critical, but rare, experience in the natural environment. Such an acquired behavior can also be modulated by the later experience as evident in the memory extinction through repetitive activation of one sensory pathway to unrelate it from another sensory experience (Robleto et al., 2004; Delamater, 2004; Bouton, 2004; Rescorla, 2004; Thanellou & Green, 2011). Critically, it remains enigmatic how the brain reconciles conflicting sensory experiences such as the coincidental and independent occurrences of stimuli to calibrate behavioral outputs. Specifically, whether excitatory and inhibitory memories develop through a common associative process or emerge as independent, parallel traces remains a fundamental question in systems neuroscience. Here we studied this issue using classical eyeblink conditioning in mice, which has been extensively studied for a century to assess learning ability and the underlying neural circuits in a variety of species including humans (Cason, 1922; Christian & Thompson, 2003; Heiney et al., 2014).

Repeated presentation of conditioned stimulus (CS; e.g., tone or light) coupled with eyelid reflex-inducing unconditioned stimulus (US; e.g., air puff to cornea) typically at a high probability (∼90%) makes animals learn to blink in response to the initially neutral CS. Such acquired responses through training are called conditioned responses (CRs). Previous studies demonstrated that the even halved pairing of sensory inputs is effective to establish CRs (Thomas & Wagner, 1964; Leonard & Theios, 1967; Gormezano & Coleman, 1975; Buchanan et al., 1997; Kehoe et al., 2008), but it remains unclear how the learning is affected by mingling of frequent neutral sensory stimulation which possibly leads to extinction. Increasing neutral CS-alone has been associated with memory erasure/extinction, and latent inhibition, the lowered efficiency of subsequent memory acquisition (Lubow & Moore, 1959; Schmajul et al., 1994; Nicholson & Freeman, 2002), although these have been regarded as distinct phenomena. Recent elegant experiments in diverse species from fly to mammals have suggested that memory extinction relies on the experience-dependent formation of another memory to counteract the original memory encoding an action (Kim et al., 2020; Felsenberg et al., 2018). However, the nature and the way of this suppressing memory to counteract the learned behavior is totally unclear in terms of cellular and circuit-level of plasticity in the mammalian cerebellar circuits. In this study, by systematically changing the enrichment of sensory pairings, we quantitatively investigated the acquisition and extinction of associative eyeblink learning. Counter-intuitively, we found that a much less or lower density of associative experience is not only sufficient but substantially more efficient per trial for the establishment of well-timed eyelid closure. Our detailed analysis of experimental data yielded a computational model demonstrating that learned behavior is established by the parallel formation of two opposing long-term memories: one for execution of eyeblink and the other for suppression. The model predicted that the suppressive memory can be pre-emptively established even before acquisition, which was experimentally validated, leading to the conclusion that the adaptive behavior is the outcome of counteraction between two opposing long-term memories.

## Results

### Eyeblink conditioning with limited paired sensory stimuli

To obtain insights into the formation and interaction of memories controlling animals’ behavior in response to distinct patterns of sensory experience, we first examined the effect of altered ratio of two sensory stimuli: tone (290 ms) and air puff (to a cornea, 40 ms) as CS and US, respectively (Fig. 1A, B). Mice were head-fixed on a treadmill during experiments (Fig. 1C) and their eyelid movements were recorded with a monochrome camera, and the eyelid closure level was quantitated by calculating the area size of an eyeball (Fig. 1D; Heiney et al., 2014; for details see Materials & Methods). Air puff delivered via a glass tube or a needle elicited quick eyeblink responses (unconditioned responses, URs), whereas tone did not affect the eyelid (Fig. 1D, before). For association of two sensory events in mice, combined and co-terminating sensory inputs of CS and US in time series are provided typically at a high ratio: 90 paired & 10 CS-alone trials (90%_CS-US_, Fig. 1A). In line with previous studies (Heiney et al., 2014; Fiocchi et al., 2022), CRs were evident in ∼10 out of 100 trials on average even on the 1st day, gradually becoming frequent (∼ 50%) and larger by ∼ day4, which remained nearly constant till day10 (Fig. 1E, F), although showing variable time courses in individual mice (Fig. S1A). On the other hand, unexpectedly, even when the pairing of CS and US was lowered to 10% (i.e., 10 paired & 90 CS-alone per day: 10%_CS-US_, Fig. 1A), mice acquired CRs during 10 days training (Fig. 1E, middle) although hitting a plateau around 30% on day6 (Fig. 1F). Thus, even 9 times less association of CS and US inputs succeeded in the establishment of eyeblink conditioning, while the final probability for the CR triggering was apparently lower with a tiny delay in reaching equilibrium (Thomas & Wagner, 1964; Leonard & Theios, 1967; Gormezano & Coleman, 1975; Buchanan et al., 1997; Kehoe et al., 1983). Notably, the number of paired trials required for CR acquisition was significantly lower in the sparse pairing groups, indicating that frequent CS-alone trials do not interfere with, but rather highlight, an underlying high-efficiency learning mechanism.

**Fig 1.**
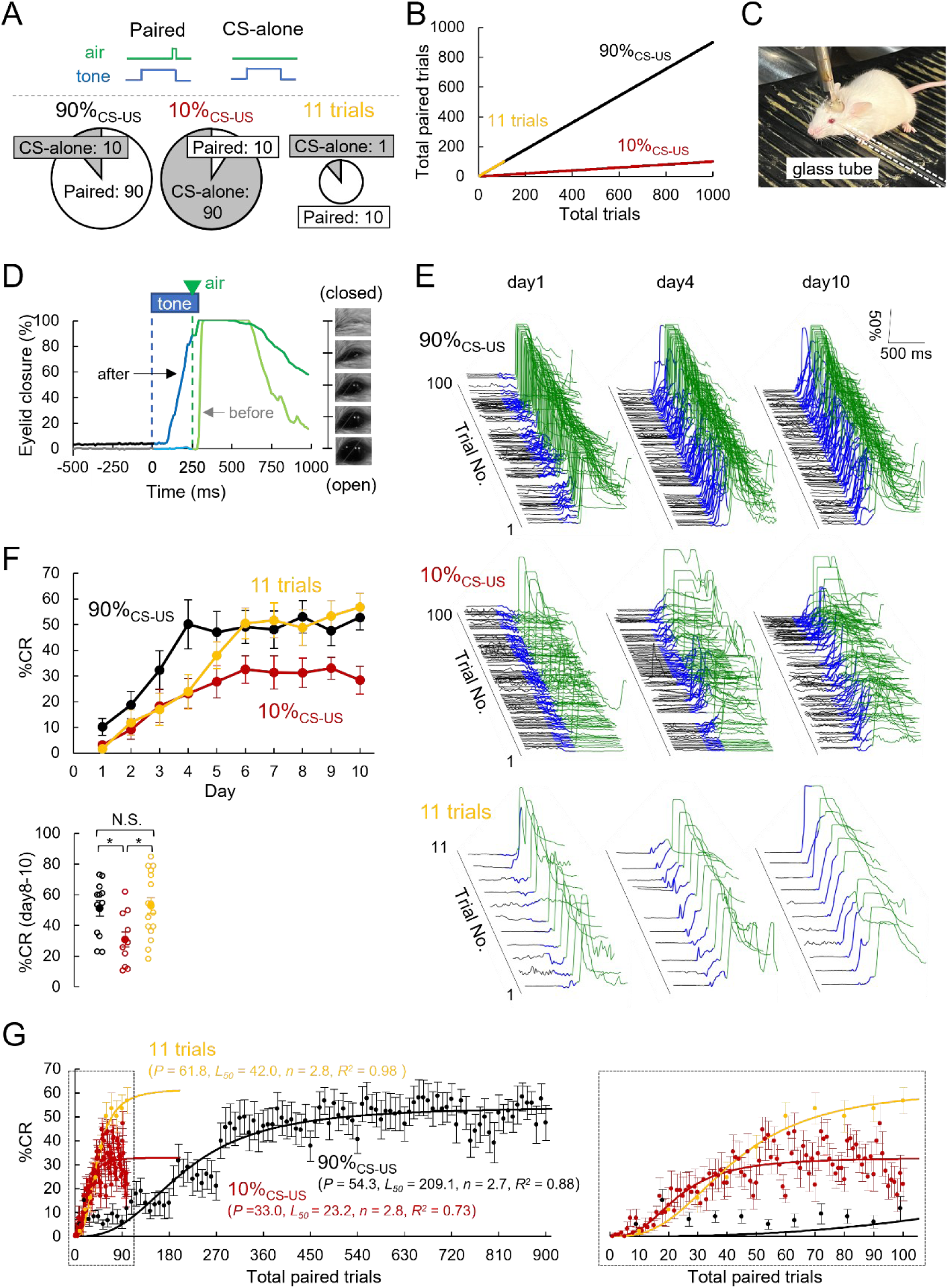
Eyeblink conditioning with distinct patterns of sensory pairings. **(A**) Schematics of CS (tone)-US (air puff) and CS-alone trials (top) and three training protocols, 90%_CS-US_, 10%_CS-US_, and 11 trials (bottom). (**B**) Total paired trials number plotted against total trials during 10 days training (90%_CS-US_, black; 10%_CS-US_, red; 11 trials, orange). (**C**) An image of a head-fixed mouse on a treadmill during experiments. A glass tube put in front of the eyelid is outlined by white dashed line. (**D**) Representative images and time courses of eyelid closure before/after the CR acquisition trainings. (**E**) Representative eyelid traces (100 or 11 trials from a single mouse) for the 90%_CS-US_ (top), 10%_CS-US_ (middle) and 11 trials (bottom) on day1 (left), 4 (middle) and 10 (right). (**F**) Daily changes of %CR established by 10 days training (top) and averaged %CR during day8-10 (bottom; 90%_CS-US_: 51.1 ± 5.0%, n = 12; 10%_CS-US_: 30.9 ± 4.9%, n = 11; 11 trials: 53.0 ± 5.1%, n = 16, *, p < 0.05 by Tukey-Kramer test), from mice showing CR establishment (CR ≥ 10% in at least 3 days). Individual, open circle; average, filled circle. (**G**) %CR averaged from each 10 or 11 trials plotted against the number of total paired trials (90%_CS-US_, n = 18; 10%_CS-US_, n = 13; 11 trials, n = 17). Fitted curves are also shown.

Two factors could underly the lower percentage of CRs (%CR) in the 10%_CS-US_ group: insufficient paired sensory experience or excessive presentation of unpaired tone. To discriminate these, another paradigm “11 trials” was examined (10 paired & 1 CS-alone per day), where the limited paired sensory experience was the same as the 10%_CS-US_, but the excessive tone-alone stimulation was omitted (Fig. 1A, B). As shown in Fig. 1E, F and S1A, even the daily 10 CS-US pairings effectively established CRs reaching ∼50% in 6 days training, similarly to the daily 100 trials in the 90%_CS-US_, although slight delay in reaching plateau. Success rate of learning in individual mice in three groups was similar, although slightly low in the 10%_CS-US_ group (Fig. S1B). These results altogether indicate that small number of paired sensory stimuli (10 pairings per day) are sufficient for the CR acquisition, while additional massive tone-alone events only lowered final probability of learned behavior. Notably, considering that the probability of air puff was just 10% out of total tone presentations in the 10%_CS-US_, the final %CR as ∼30% could be rather regarded as exaggerated occurrence of acquired behavior.

While the 10% CS-US group exhibited a slower increase in daily %CR compared to the 90% group, a strikingly different picture emerged when the data were normalized by the number of paired trials. When the averaged %CR was plotted as a function of total CS-US pairings, surprisingly, the 10%_CS-US_ and the 11 trials groups exhibited very quick establishment of CRs compared to the conventional 90%_CS-US_ group (Fig. 1G). The courses of increase in %CR were fitted by the following equation:

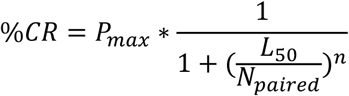

, where *P_max_*, *L_50_*, *n*, and *N_paired_* represent the maximal %CR, the number of paired stimuli for the half-maximal learning, the steepness of learning, and the number of given paired trials. In sharp contrast to the similar *n* in three groups, *L_50_* reflecting the number of sensory inputs needed for learning was ∼6 folds higher in the 90%_CS-US_ group. Thus, the sparse pairing protocol achieved superior learning efficiency, reaching the *L_50_* with nearly six-fold fewer pairings, in contrast to previous findings in rabbits (Thomas & Wagner, 1964; Leonard & Theios, 1967; Gormezano & Coleman, 1975). Step-by-step pattern of %CR increases were evident in the 90%_CS-US_ group during initial 4 days, implying some daily upper limit for learning such as a gradual attenuation of its efficiency during repetitive training (Tait et al., 1983; Kehoe & White, 2002). Intriguingly, the %CR in the 10%CS-US group was initially compatible at the beginning of each daily trials to the 90%CS-US and the 11 trials groups, but selectively exhibited a gradual decline during daily sessions (Fig. 1G, inset). Thus, tone-alone trials do not disturb the acquisition, but distract the expression of learned CRs. Furthermore, taking into consideration that the total trial number is identical in 10%- and 90% CS-US groups, the “within-session decline” of %CR evident only in the 10% group implies a gradually emerging process to actively suppress the CR occurrence.

The acquired eyeblink responses were similarly sophisticated in timing and amplitude although the averaged eyelid closures in the 10%_CS-US_ group were apparently smaller because of larger variability of peak timing (Fig. 2A-C). The CR amplitude grew up gradually during days of training and showed faster saturation in the 10%_CS-US_ and the 11 trials groups in relation to the number of paired trials, while onset/peak timings got quickly optimized in all groups (Fig. S2A-E) (Heiney et al., 2014; Kehoe et al., 2008; Fiocchi et al., 2022; Balsam et al., 2002; Ohyama & Mauk, 2001; Kehoe et al., 2014). With a closer look, the 10%_CS-US_ group occasionally exhibited rapid and <u>c</u>omplete <u>e</u>yelid <u>c</u>losure (RCEC), reaching 98% eyelid closure during 80-180 ms from the tone onset, which looked similar to that of spontaneous eye blinking (Fig. S3A-C). As a result of more RCEC occurrence, well-timed blinking relative to the timing of air puff, which was regarded to be cerebellum-dependent CRs, was slightly rarer in the 10%_CS-US_ group (Fig. S3D), while total probability of eyeblink responses including CRs and RCEC was almost compatible in three groups (Fig. S3E).

**Fig 2.**
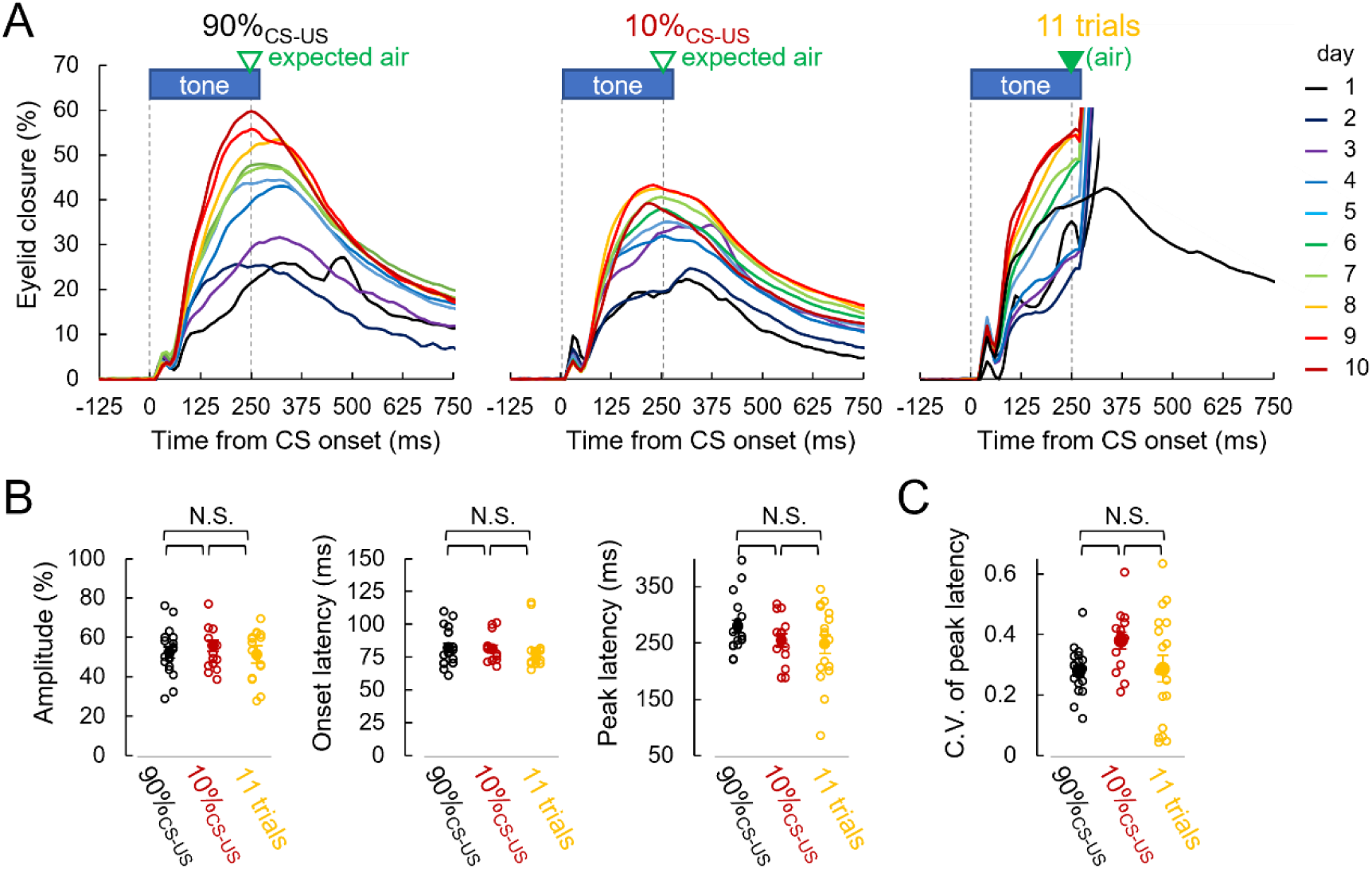
Learned eyelid responses. **(A**) Time courses of averaged eyelid closure for each day (90%_CS-US_: n = 18 obtained from CS-alone trials; 10%_CS-US_: n = 13 from CS-alone trials; 11 trials: n = 17 from paired & CS-alone trials). For 11 trials, a black trace obtained from averaging eyelid closures in CS-alone trials during 10 days is also shown. For averaging, only eyelid traces showing responses were used. Baseline level before CS was aligned to 0% for each. (**B**, **C**) Averaged amplitude (left), onset latency (middle) and peak latency (right) of eyeblink responses (B), and coefficient of variance for peak latency (C) obtained from all trials with responses. Individual, open circle; average, filled circle. Tukey-Kramer test.

### Critical role of DCN in CR expression

Deep cerebellar nuclei (DCN) plays an important role in expression of CRs in rabbits (Christian & Thompson, 2003; McCormick & Thompson, 1984; Yeo et al., 1985) and mice (Heiney et al., 2014; Sakamoto & Endo, 2008). To examine whether the CRs acquired with 10%_CS-US_ also rely on DCN, we pharmacologically inhibited neuronal activity by infusing muscimol, a selective GABA_A_ receptor agonist, to the anterior interpositus part of DCN through a cannula implanted in the already CR-acquired mice (Fig. 3A). As shown in Fig. 3B, cannula position and estimated area for diffusion of muscimol were similar in the two groups (90% and 10% CS-US pairing groups). Muscimol administration (evaluated by at least 30 trials given before and after its infusion) potently abolished the CR expression in both the 90% and 10% CS-US pairing groups, followed by almost complete recovery on the next day (Fig. 3C). Averaged traces for eyelid closure indicated marked abolishment of CRs by muscimol, but not by artificial cerebrospinal fluid (aCSF), without affecting the URs upon air puff (Fig 3C-F). Taken these results together, it was revealed that CR is expressed depending on the DCN activity irrespective of the altered enrichment of paired sensory experience.

**Fig 3.**
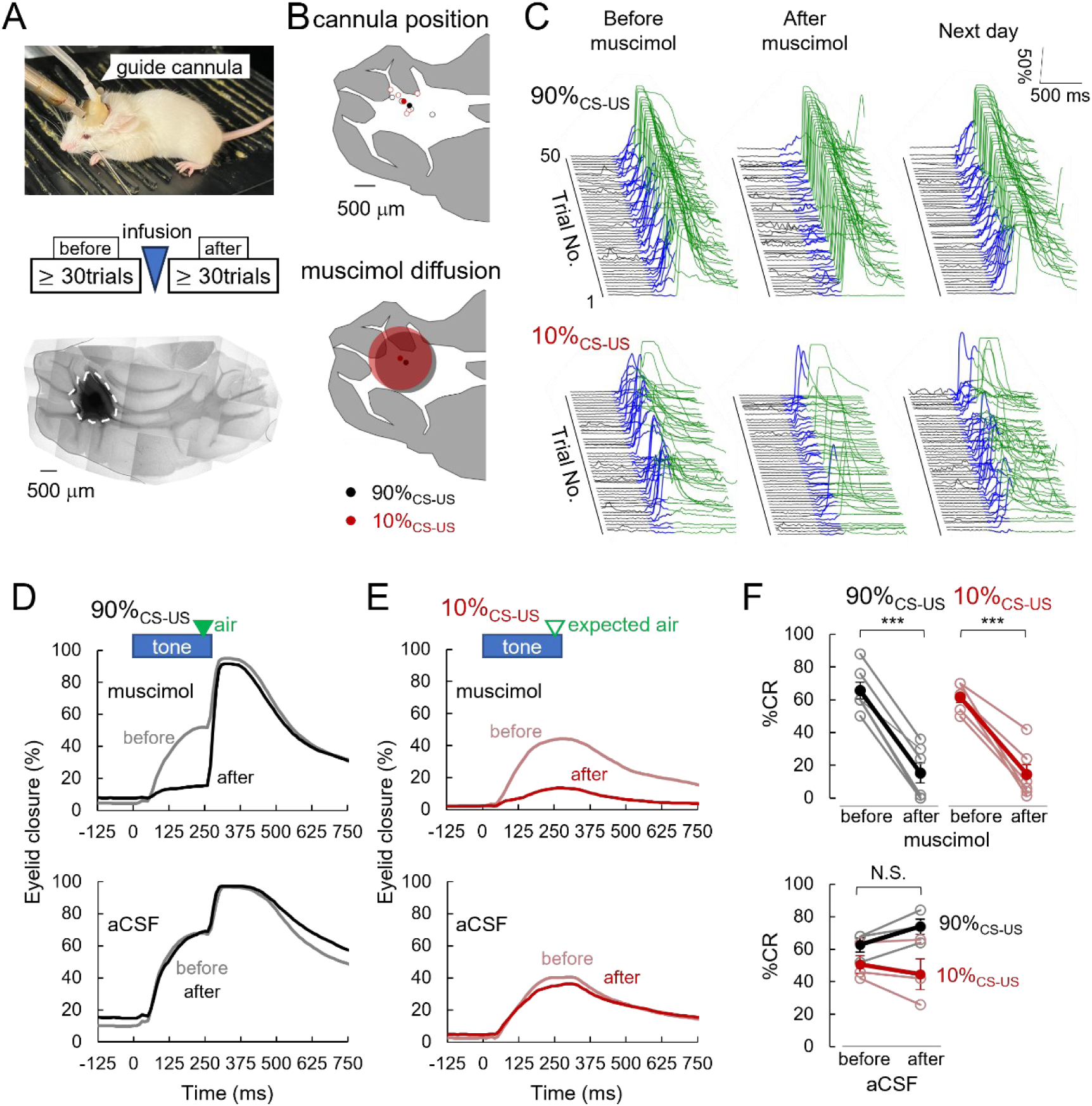
Critical role of DCN activity for occurrence of established CRs. **(A**, **B)** Schematics of pharmacological inactivation by muscimol (A) and the target location (B) in the cerebellum. Muscimol or aCSF was infused through an implanted guide cannula (top). Mice had at least 30 trials before/after muscimol or aCSF infusion (middle). Area for muscimol diffusion identified from fluorescence on a horizontal cryosection is shown in A, bottom (white dashed line), and summarized in B (90%_CS-US_, n = 5; 10%_CS-US,_ n = 5). Cannula tip positions are also shown in B, top (90%_CS-US,_ n = 4; 10%_CS-US,_ n = 5). The distance of each average cannula position was less than 150 μm and the diameter of muscimol diffused area was 1.45 ± 0.19 mm for 90%_CS-US_; 1.50 ± 0.12 mm for 10%_CS-US_. (**C**) Representative eyelid traces from a single mouse in 90%_CS-US_ (top) and 10%_CS-US_ (bottom) before, after and the next day of muscimol infusion. (**D**, **E**) Averaged eyelid traces before/after muscimol (top) or aCSF (bottom) infusion in 90%_CS-US_ (D, n = 6) and 10%_CS-US_ (E, n = 6). Eyelid traces from all acceptable trials were used for averaging. (**F**) %CR changes before/after infusion of muscimol (top) and aCSF (bottom). ***, p < 0.001 by paired t-test.

Scatter plots of averaged eyelid closure amplitude against %CR exhibited a tendency of the 90%_CS-US_ showing positive relation between them (Fig. S4A, left), which was preserved even after the DCN inactivation. In contrast, the 10%_CS-US_ group showed less correlation between %CR and eyelid closure amplitude, sometimes exhibiting unaltered amplitude even after the DCN inactivation (Fig. S4A, right). Furthermore, RCEC sometimes evident in the 10%_CS-US_ group tended to resist to the DCN inactivation (Fig. S4B). Thus, limited experience and disturbance by lots of CS-alone inputs in the 10%_CS-US_ group might be partly compensated by recruiting neural circuits other than the cerebellar circuit for eliciting eyelid closure (Kaminer et al., 2011; Kaminer et al., 2015; Bologna et al, 2024). Nevertheless, the similar impact of DCN inactivation on CR execution, irrespective of altered enrichment of sensory pairings, indicates that the occurrence of learned behavior is primarily governed by the DCN, probably with slight difference in experience-dependent circuits modulation mainly in the cerebellum.

### Distinct extinction dynamics of CRs acquired with altered enrichment of sensory experience

In order to evaluate the qualitative identity of CR-memories acquired by different patterns of experience, we examined memory-extinction paradigm. Following 10 days of acquisition training, 100 tone-alone (i.e. 100 CS-alone) trials were daily applied for 4 days. CR occurrence gradually decreased during the extinction session with clearly different kinetics according to the distinct pattern of acquisition (Fig. 4A-C). Looking at gradual changes during daily sessions, %CR substantially recovered at the beginning of individual daily sessions (Fig. 4B), known as spontaneous recovery (Robleto et al., 2004; Delamater, 2004; Bouton, 2004; Rescorla, 2004; Thanellou & Green, 2011). The CR extinction profiles (Fig. 4C) could be fitted to single exponential curves, as characterized by the following equation:

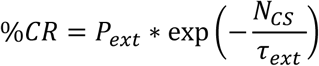

, where *P_ext_*, *τ_ext_* and *N_CS_* represent the initial value of %CR, the factor for resistance of CR to be extinct, and the number of CS-alone trials during each day, respectively. The CRs established by rich paired stimuli seem to exhibit ∼4 times greater resistance to extinction in mice, which is distinct from humans showing higher resistance in partial reinforcement-acquired CRs (Grant & Schipper, 1952; Humphreys, 1939; Grant & Hake, 1951; Moore & Gormezano, 1963) and also rabbits with almost similar resistance irrespective of sensory enrichment during acquisition (Thomas & Wagner, 1964; Gormezano & Coleman, 1975). In clear contrast to the %CR attenuation (i.e., reduction in percentage of occurrence), eyelid closure amplitude, if occurred, was kept high during the extinction sessions in all groups, although the 11 trials group showed slight decline (Fig. 4D, E). Furthermore, interestingly, eye blinking like spontaneous responses tended to be frequently observed during extinction (Fig. S3C, 4F, G). As for the timing control, response onset showed delay during extinction although peak latency exhibited large variabilities among individuals (Fig. 4H), as previously reported (Kehoe et al., 2014; Medina et al., 2002). Taken together, eyeblink extinction learning is expressed as purely decreased probability, without changing the behavior dynamics, of blinking in response to the repetitive tone-alone experience, implying the preserved memory for well-timed blinking in the neuronal circuit, in line with the idea that the CR extinction is mediated by formation of new memory to counteract the CR occurrence (Kim et al., 2020).

**Fig 4.**
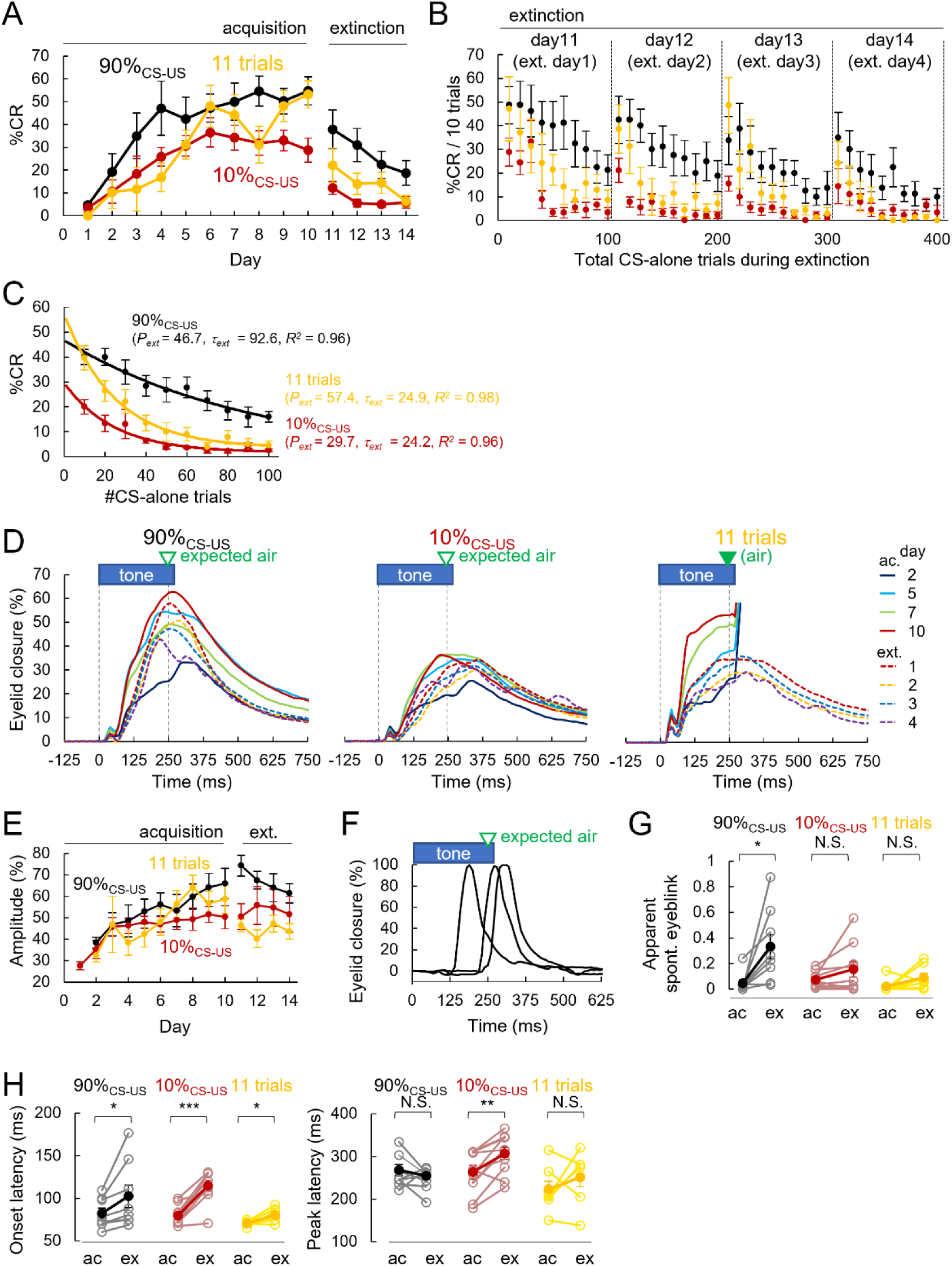
Dynamics of CR probability and eyelid closures during memory extinction. **(A**, **B**) Daily %CR changes during 14 days of training consisting of acquisition and extinction sessions (A) and %CR averaged from each 10 trial during 4 days extinction (B). 90%_CS-US_, n = 8; 10%_CS-US_, n = 9; 11 trials, n = 7. (**C**) Averaged %CR changes of every 10 trials from the beginning of extinction day1-4 were plotted against the number of CS-alone trials with fitting to single exponential curves. (**D**, **E)** Averaged eyelid closure traces on acquisition day2, 5, 7, 10 and extinction day1-4 (D), indicated by different colors, and daily changes of averaged amplitude of eyelid closure (E) during acquisition and extinction paradigms. For 90%_CS-US_ and 10%_CS-US_, from CS-alone trials; for 11 trials, from both paired & CS-alone trials. The baseline level before CS was aligned to 0% for each. (**F**) Example eyelid traces for responses like spontaneous eyeblink (> 90% of eyelid closure with half-width < 150 ms). (**G**) Ratio of spontaneous eyeblink-like ones to all responses. Acquisition, ac; extinction, ex. *, p < 0.05 by paired t-test. (**H**) Onset latency (left) and peak latency (right) during acquisition and extinction training. *, p < 0.05; **, p < 0.01; ***, p < 0.001 by paired t-test.

### Modeling CR acquisition and extinction through integrating total history of sensory inputs

To shed light on the underlying mechanisms of distinct dynamics of CR acquisition and extinction with altered pairings of sensory stimuli, we attempted to develop a model that explains our observations (see Figs. 1 and 4). Several lines of evidences in our experiments, such as the apparent differences in rate of acquisition, saturated level, but preserved eyeblink dynamics and the daily spontaneous recovery during extinction (see Fig. 4), indicated interaction of independently established two memory components for CR, one encoding execution and the other suppression, as previously suggested (Robleto et al., 2004; Delamater, 2004; Bouton, 2004; Rescorla, 2004; Felsenberg et al., 2018). Thus, we built a model assuming parallel formation of counteracting memories for CRs, as follows (here, in modeling section, we denote %CR as *CR_ratio_*):

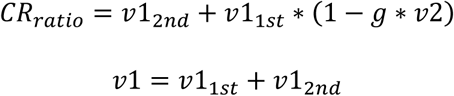

, where *v1* is the memory for CR execution consisting of two components, *v1_1st_* and *v1_2nd_* with distinct susceptivity to the CR suppressing memory, *v2*. *g* represents a gate for *v2*, increasing as a function of total amount of CS inputs (i.e. total trial number) (Fig. 5A; for details see Materials and Methods). The larger *v1* results in higher *CR_ratio_*, while the larger *v2* leads to smaller *CR_ratio_*. *v1_1st_*, *v1_2nd_* and *v2* were assumed to build up trial by trial based on the Rescorla and Wagner model (Rescorla & Wagner, 1972) with some modification, defined as follows:

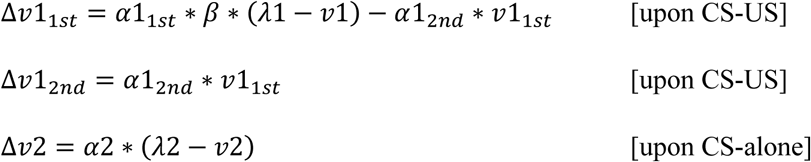

, where *α1_1st_, α1_2nd_* and *α2* represent rate constants for formation of memories *v1_1st_, v1_2nd_* and *v2*, respectively. *v1_1st_* was assumed to be formed upon the CS-US pairings at a rate *α1_1st_* multiplied by an efficiency factor *β*, and to be gradually turned into the extinction-resistant *v1_2nd_* at a rate *α1_2nd_*. Stronger resistance of CR to extinction in the 90%_CS-US_ was made possible by this scheme (see Fig. 4C). Daily upper limit of learning, inferred from Fig. 1G, was implemented in the model by gradual decline of the *α1_1st_* efficiency factor *β* as total paired trials increase within a session (Fig. S5A).

**Fig 5.**
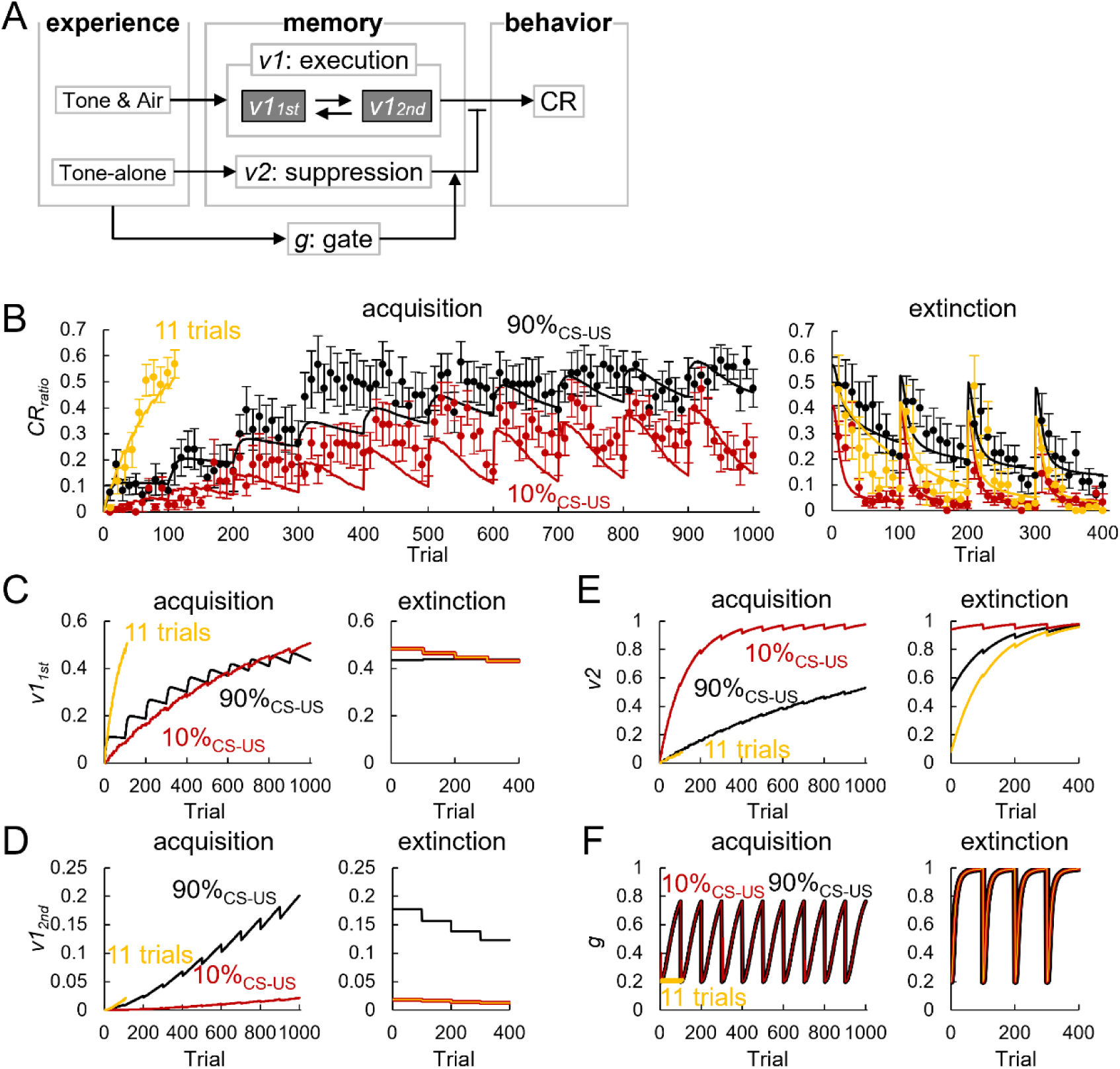
Modeling of formation of counteracting memories for CRs. **(A)** Model scheme for input-dependent memory formation for CR execution and its suppression. **(B)** Simulated *CR_ratio_* plotted as lines against total trial number during acquisition (left) and extinction (right) paradigms, together with %CR averaged from each 10 or 11 trial recorded in experiments (dots, 90%_CS-US_, n = 12; 10%_CS-US_, n = 11; 11 trials, n = 16). (**C-F**) Two components for memory to execute CR, *v1_1st_* (C) and *v1_2nd_* (D), the counteracting memory to suppress CR, *v2* (E), and its gate *g* (F) are plotted against total trial number during acquisition (left) and extinction (right) in simulation.

Both the acquisition and extinction processes of *CR_ratio_* were simulated in the model, and distinct dynamics for three experimentally-tested patterns were nicely reproduced (Fig. 5B-F). The 90%_CS-US_ group hit daily plateau of CR acquisition, whereas the 11 trials group showed more efficient learning as observed in experiments. In the 10%_CS-US_ group, even after reaching apparent saturation, *CR_ratio_* exhibited daily quick increase and following “within-session decline” because of gradual gating of slowly summating *v2* to counteract *v1_1s_*_t_ via activation of *g* by repeated trials (Fig. 5B, E, F).

We also conceived another model in which the suppressing memory, *v2* was short-lived and refreshed by the next session, rather than long-lasting (compare Fig. S6B with Fig. 5E). As shown in Fig. S6A, this model also succeeded in reproducing experimental observations. In this version, notably, daily spontaneous recovery during extinction sessions can be explained just by the complete refreshment of *v2* irrespective of the inclusion of a gating mechanism of *v2* (Fig. S6B, C). However, the “within-session decline” during acquisition in the 10%_CS-US_ was still strongly dependent on the gating mechanism: eliminating the gate made the “within-session decline” more rapid and potent than actual experimental data (Fig. S6C). Thus, in any cases, the memory *v2* to suppress CRs are to be adequately controlled by the online experience of sensory inputs, as has been suggested in a previous study showing that the decline of %CR speeded down with occasional paired sensory inputs during the extinction session (Gibbs et al., 1978).

### Model-driven discovery of prior-formed suppression memory

To evaluate the qualitative difference of learning dynamics generated by two *v2,* characterized with distinct temporal lifetime, long- or short-lasting one, we attempted to obtain a prediction from the two models. Repetitive CS-alone trials presented for days in prior suppressed the later *CR_ratio_* increase in the model with long-lasting *v2*, but not in the model assuming short-lasting *v2* (Fig. 6A). Thus, the modeling analysis yielded a prediction: if long-lasting, the suppressing memory *v2* can be formed in prior to the start of CR acquisition. This prediction was tested by behavioral experiments presenting 100 CS-alone trials for 4 days prior to the start of 90% CS-US pairing paradigm (the Pre group). Strikingly consistent with the model prediction, as shown in Fig. 6A and B, the Pre CS-alone trials potently inhibited the initial growth of CRs, which has also been known as latent inhibition (Lubow & Moore, 1959; Schmajuk et al., 1994; Nicholson & Freeman, 2002), without affecting the dynamics of acquired eyelid closure (Fig. 6C). Furthermore, despite the later training paradigm by itself was the 90% pairing of CS and US, surprisingly, the increase of %CR in the Pre-group showed quite similar pattern to that in the 10%_CS-US_ groups rather than the 90%_CS-US_ (Fig. 6B), characterized by “within-session decline” (Fig. 6D).

**Fig 6.**
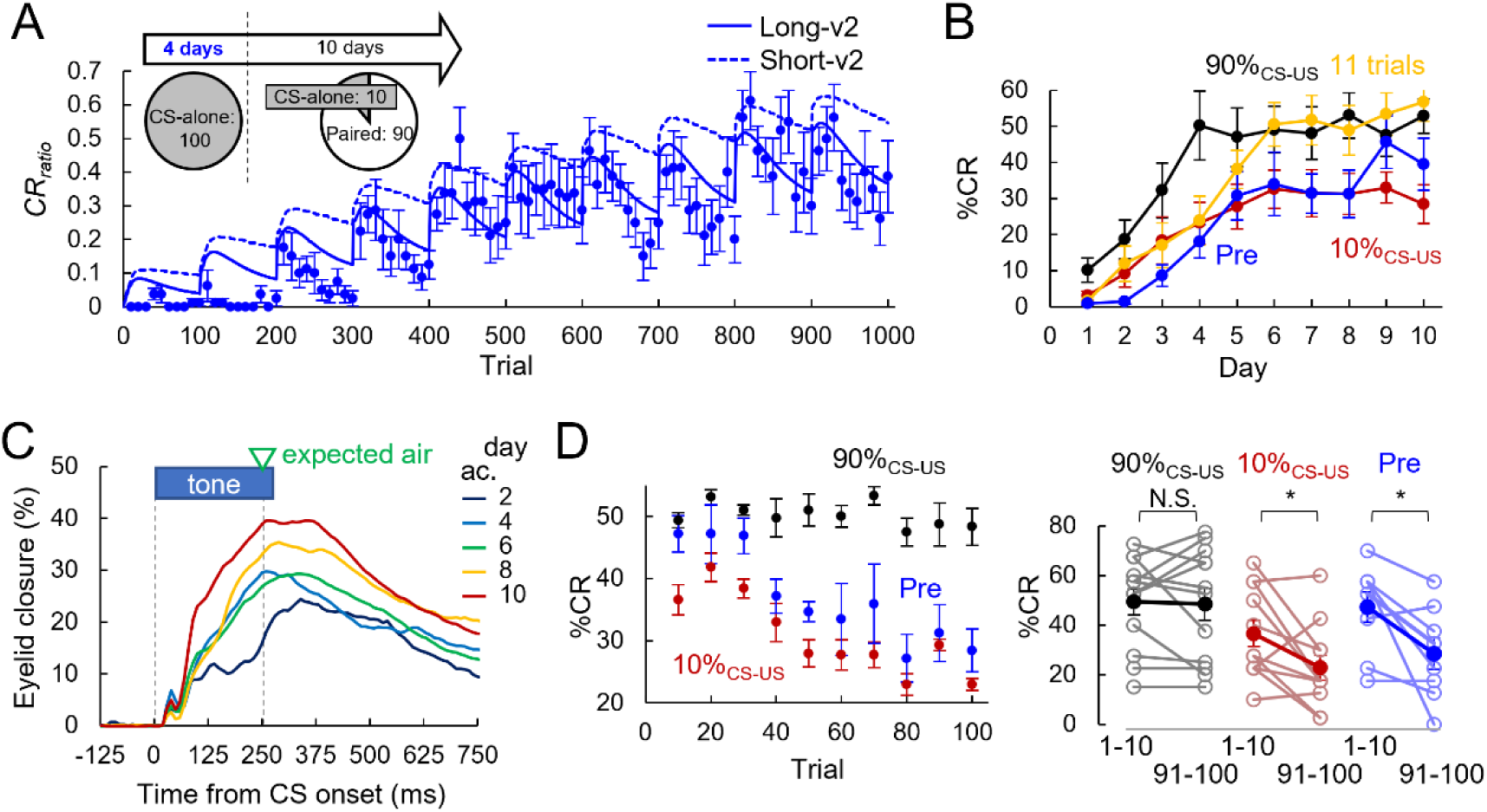
Prediction and validation of long- or short-lasting suppression memory model. **(A)** Prediction of *CR_ratio_* changes during acquisition by 90%_CS-US_ following the 4 days CS-alone exposure (Pre) in models assuming long (solid line)- or short (dotted line)-lasting *v2*. Actual experimental results of CR ratio averaged from each 10 trial (n = 8, dot) are also shown to evaluate the model prediction. Training protocols for the Pre group is also shown as inset. **(B**) Daily %CR changes during 10 days of acquisition training. Pre, n = 8. Data for the 90%_CS-US_ (n = 12), 10%_CS-US_ (n = 11), and 11 trials (n = 16) shown in Fig. 1F are also superimposed for comparison. **(C**) Averaged eyelid closures observed in CS-alone trials on day2, 4, 6, 8, 10 in the Pre group mice which were given CS-alone trials for 4 days in prior to the 90% CS-US pairing. (**D**) Averaged %CR for every 10 trials on day7-10 were plotted against trials number (left). Comparison of averaged %CR in 1-10 and 91-100 trials (right). *, p < 0.05 by paired t-test.

Accordingly, the zig-zag pattern in *CR_ratio_* dynamics was more evident in the model with long-lasting *v2* (Fig. 6A). Thus, the prediction of model assuming long-lasting v2 was validated by the animal behavior experiments, and amazingly it was revealed that even before the acquisition of CRs, the prior experience of CS-alone inputs forms negative counteracting memory for not only the later acquisition but also the execution probability of CRs after acquisition, as if the CR memory was established through the limited pairings. Collectively, we conclude that the association of CS and US forms a memory to execute CRs, while CS-alone experience forms a counteracting long-term memory to suppress CRs in parallel and independently from the situation of the target positive memory.

## Discussion

Changing the training paradigm of cerebellum-dependent eyeblink conditioning, here we demonstrated that the learning efficiency is paradoxically facilitated by restricting the number of daily trainings. Distinct occurrence profiles of the learned behavior without changes in blinking dynamics during the acquisition and extinction highlighted the sensory experience-dependent parallel formation of long-term memories for the CR execution and its suppression. The different counteraction of the two memories explained patterns of acquired behavior occurrence with distinct sensory experience in acquisition, extinction, and even latent inhibition. Thus, our present study provides a previously unprecedented simple framework for learning assuming only two memories, by which distinct sensory experiences in combination and timeline can be integrated in animals brain.

### CR acquisition with partial reinforcement

Eyeblink conditioning is conserved widely in mammals, showing specie-dependent different relations between the training patterns and the learning dynamics. Humans show %CR closely related to the ratio of pairings (Grant & Schipper, 1952; Grant & Hake, 1951; Moore & Gormezano, 1963; Ross, 1959), while rabbits supra-linearly exhibit CRs upon training with smaller possibility of sensory pairings (Thomas & Wagner, 1964; Leonard & Theios, 1967; Gormezano & Coleman, 1975; Buchanan et al., 1997). As demonstrated here, even much smaller 10% pairing is sufficient for the CR acquisition in mice (see Fig. 1F), showing rather inversed relationship between the enrichment of associative experience and the learning efficiency (see Fig. 1G), which contrasts to the previous findings in rabbits typically showing equal or less efficiency with partial reinforcement (Thomas & Wagner, 1964; Leonard & Theios, 1967; Gormezano & Coleman, 1975). Apparent slower growth of %CR in mice than in rabbits during days of trainings (Thomas & Wagner, 1964; Heiney et al., 2014; Leonard & Theios, 1967; Gormezano & Coleman, 1975; Fiocchi et al., 2022), implies more potent upper limit of learning during a certain period in mice, spoiling surplus reinforcement experience. This could be responsible for the apparent higher learning efficiency upon limited reinforcement. Indeed, learning efficiency can be estimated higher also in rabbits with smaller density of training, when compared with a training paradigm accompanied with daily saturation or decline of %CR (Tait et al., 1983; Kehoe & White, 2002).

### Candidates for neural circuits generating distinct profiles in CR execution

We demonstrated a critical role of cerebellar circuit in expression of learned eyeblink irrespective of the enrichment of reinforcement (Fig. 3; Christian & Thompson, 2003; Yeo et al., 1985; McCormick et al., 1982; Krupa et al., 1993). Long-term depression (LTD) at parallel fiber (PF) synapses on a Purkinje cell (PC) has been focused as a substrate of memory formation for CRs (Koekkoek et al., 2003). In addition, plasticity at mossy fiber (MF) synapses on neurons in DCN also plays a role, contributing to higher resistance to extinction presumably requiring more experience for its establishment compared to PF-PC LTD (Garcia & Mauk, 1998; Medina et al., 2001; Ohyama et al., 2006; Antonietti et al., 2016; Antonietti et al., 2017; Broersen et al., 2023). Given these findings, the PF-PC LTD and plasticity at MF-DCN synapses might correspond to the extinction-sensitive (*v1_1st_*) and -resistant (*v1_2nd_*) memories for CRs in our hypothetical model (see Fig. 5A).

While classical Rescorla-Wagner models account for the net associative strength, our findings necessitate a model with dual, independent components to explain the dissociable kinetics of CR acquisition and its within-session suppression, as previously suggested as independent memory formation for extinction of learned behavior (Robleto et al, 2004; Delamater, 2004; Bouton, 2004; Rescorla, 2004; Felsenberg et al., 2018). Accordingly, our experiments and modeling together highlighted an importance of the memory counteracting CR execution, not only for the extinction, but also for the fine-tuned probability to execute learned eyelid closure. In addition, surprisingly, our data suggest that the inhibitory memory trace (*v2*) can be somehow long-lasting and formed even prior to the formation of positive memory to execute eyeblink. Crucially, our model’s prediction that the inhibitory memory trace can be surreptitiously established even in the absence of behavioral CR was empirically validated. Thus, *v2* seems to practically underlie the lowered efficiency in acquisition after the prior CS presentations, known as latent inhibition (Lubow & Moore, 1959; Schmajul et al., 1994; Nicholson & Freeman, 2002). Recently, it was demonstrated in *Drosophila* that odor pre-exposure forms an independent aversive memory that competes with subsequent learning, providing a mechanistic basis for latent inhibition and also extinction learning (Jacob et al., 2021). Our results extend this concept to the mammalian cerebellum, demonstrating that such “pre-emptive” inhibitory memories are not only formed prior to acquisition but also dynamically interact with excitatory traces to fine-tune motor outputs on a trial-by-trial basis.

One candidate mechanism of the inhibitory memory trace in the cerebellum is long-term potentiation (LTP) between PF and PC in cerebellum (Kim et al., 2020; De Zeeuw, 2021; Kostadinov & Häusser, 2022). Cerebellar extinction was shown to involve a reversal of synaptic plasticity, such as the transition from LTD to LTP at PF-PC synapses, driven by the deprivation of climbing fiber signals (Medina et al., 2002). Our present study presented another suppressing process for the memory under conditions of sparse reinforcement: independent inhibitory memory (*v2*) tunes the behavior while keeping robust excitatory memories (*v1*). It is conceivable that LTD and LTP at PF-PC synapses are formed at distinct micro-zones (De Zeeuw, 2021; Kostadinov & Häusser, 2022), and outputs from these PCs counteract at identical DCN neurons. This bidirectional control of learning gain resonates with the cellular mechanisms proposed for vestibulo-ocular reflex (VOR) adaptation, where the balance between LTD and LTP at PF-PC synapses mediates the up- and down-regulation of motor gain (Raymond and Lisberger, 1998; Boyden and Raymond, 2003). While VOR adaptation is often explained by the shifting balance of these opposing plasticities within a single pathway, our findings in eyeblink conditioning extend this concept by demonstrating that ‘gain’ can also be regulated through the parallel accumulation of an independent inhibitory memory. In our model, the ‘10% reinforcement’ advantage is not merely a result of less synaptic saturation, but an manifestation of an optimized gain control where the inhibitory memory (*v2*) prevents excessive responding without erasing the excitatory trace. This suggests a universal principle in cerebellar computation: whether through bidirectional plasticity at a single synapse or through the parallel recruitment of opposing memory pathways, the cerebellum actively calibrates its output gain to match the statistical structure of environmental feedback. Alternatively, the suppression of CR-executing processing might be enabled in the cerebellar circuit by dynamic plasticity at GABAergic synapses, for example between molecular layer interneurons and PCs (Kawaguchi & Hirano, 2000) and/or those between PCs and neurons in DCN (Aizenman et al., 1998). In addition, outside of cerebellum, CR extinction-related hippocampus (Schmaltz & Theios, 1972; Akase et al., 1989), or dopaminergic signals from ventral tegmental area (VTA) to nucleus accumbens (Kutlu et al., 2022) and posterior part of striatum (Menegas et al., 2017) related to latent inhibition might affect plasticity at various synapses in the cerebellar cortex and nuclei.

Our present data demonstrated that negative effects on CR execution in the latent inhibition, extinction, and “within-session decline”, could be integrated into the sole inhibitory memory trace *v2* in the simple framework (see Fig. 5A), which corresponds to mechanistic clarification of the nature of a century debate on the latent inhibition, primarily explained as the lowered attention. Gating of such an inhibitory memory across past, online, and ongoing sensory events highlights that the cerebellum constantly integrates sensory history to calibrate future motor outputs. Taken all these findings together, by shedding light on the counteractions of opposite memories integrating distinct sensory inputs, the present study opens a door to clarify the key neuronal mechanism to fine-tune the learned behavior reflecting the whole history of experience, not only for eyeblink conditioning but also for other associative learning paradigms like fear conditioning and reward-related reinforced learning.

## Materials & Methods

### Animal usages

Adult male FVB/NJcl mice, ≥ 8 weeks old, were used in all experiments (n = 86). Mice were housed singly with *ad libtum* access to food and water in a room with 12 hours of light/dark cycle. All animal procedures were performed by the same experimenter in accordance with guidelines on animal experimentation in Kyoto University and approved by the local committee in Graduate School of Science, Kyoto University.

### Surgery

Custom-made aluminum head-plates were mounted on mice before experiments. Mice were anesthetized with intraperitoneal injection (0.2-0.3 ml/mouse) of mixture of 0.9% ketamine (Daiichisankyo) and 0.2% xylazine (Bayer). The scalp was cut off to expose the skull and mice were stereotaxically fixed on an operating table (Narishige). The underlying membranous tissue was removed with cotton swabs and small holes were made around bregma using a drill. Screws were loosely tightened to the holes. The head plate was placed on the skull surface and fixed to the screws and the skull with dental cement (UNIFAST Trad, GC). Mice had at least 30 hours of recovery period before experiments started.

To pharmacologically inactivate DCN, a guide cannula (RWD Life Science) was implanted for some mice (n = 12) together with a head plate (see Fig. 3A). A small hole (∼6.0 mm posterior, ∼2.0 mm lateral from bregma; Paxinos & Franklin, 2008) was drilled and the cannula tip was approached to the brain surface using a stereotaxic micromanipulator. The cannula was inserted 1.8-2.3 mm vertically and fixed with dental cement. An internal dummy cannula was inserted to the guide cannula to prevent clogging.

### Eyeblink conditioning, acquisition and extinction

The basic apparatus was composed of a self-made treadmill on which mice were head-fixed during experiments, a camera (CS135MU, Thorlabs) for eyelid closure recording, a speaker for tone stimuli, a solenoid for air stimuli, and a PC (XPS8940, Dell) for recording and stimuli control (Fig. S7A; Heiney et al., 2014). The treadmill was composed of cylindrical rotating rods and rubber sheet. The implanted head-plate was fixed to a manipulator (SM-19, Narishige) ∼2 cm above the treadmill with a screw (Fig. 1C). The camera was mounted on a metal rod to monitor a left eye. Mice were illuminated by white LED light and eyelid movements were recorded with 100 frames/sec. The speaker was placed back left side of mice. A gas cylinder was connected to the solenoid and air was delivered to a left cornea when the solenoid opened. The camera, speaker and solenoid were controlled in timing with Python through an AD/DA converter (DIO-8/8B-UBT, Y2 Corporation).

Mice were randomly divided into three groups with distinct trainings (90%_CS-US_, n = 22; 10%_CS-US_, n = 20; 11 trials, n = 22), as follows (see Fig. 1A): 90%_CS-US_ group, 10 blocks of 10 trials per day, with each block composed of 9 paired & 1 CS-alone trials; 10%_CS-US_ group, 10 blocks of 1 paired & 9 CS-alone trials; 11 trials group, 10 paired & 1 CS-alone trials per day. Each mouse had at least 6 days of training. Mice had habituation period without recording and stimuli delivery for ≥ 15minutes on the treadmill before training each day. The CS (tone, 3.5 kHz, 75 dB, 290 ms) was co-terminated with the US (air puff, 15-30 psi, 40 ms) in CS-US paired trials; on the other hand, the tone CS was given without the US presentation in CS-alone trials (Fig. 1A). The US was delivered via a glass tube (i.d. = 0.5 mm) or a 21 gauge needle placed ∼2 mm away from a mouse cornea (Fig. 1C) and given in an almost identical manner through experiments except for gradual increase in pressure as days of training proceeded (Fig. S7B). The order of paired and CS-only trials was randomly determined in each block. The inter-trial interval was set to 10-24 sec randomly for each trial.

Some mice (90%_CS-US_, n = 8; 10%_CS-US_, n = 9; 11 trials, n = 7) had 4 consecutive days of extinction paradigm, 100 CS-alone trials/day, following the 10 consecutive days of acquisition. Other mice (n = 8) had exposure to daily 100 CS-alone trials for 4 consecutive days prior to the 90%_CS-US_ trainings (Pre group). When training was accidentally interrupted in the first block because of system’ s error, although very rare, daily traning was started over again.

### Data analysis

Recorded videos for eyelid movements were converted into numerical trace by calculating the area of an eyeball in individual frames (Fig. 1D; Heiney et al., 2014). Full eyelid open/close corresponded to 0/100% respectively. Reflection of light and infiltration of whisker were approximately calibrated. Trials were omitted from analysis when significant activities other than eyelid closures were observed, or recording or saving of eyelid movements was not performed properly. Eyelid closure amplitude was obtained by subtraction of pre-CS baseline from the maximal eyelid closure during 80-250 ms after the CS onset (for CS-US paired trials) or 80-470 ms (for CS-alone trials). When the amplitude was ≥ 15%, this trial was regarded as one with a response and further analysis was conducted. Onset of eyelid closure was visually identified. The *⍺*-startle, a startle reflex to tone, was carefully removed from the analysis by visual identification. The responses with averaged eyelid closure amplitude during 220-250 ms ≥ 15% and larger than that during 120-150 ms, were regarded as CRs.

### Pharmacological inactivation of DCN neurons

DCN neurons in CR-learned mice (90%_CS-US_, n = 6; 10%_CS-US_, n = 6) were inactivated by muscimol (1 or 5 mM, Nakalai Tesque) administration through an implanted guide cannula (see Surgery). Muscimol was diluted in aCSF (NaCl, 125 mM, NaHCO_3_, 25 mM, NaH_2_PO_4_, 1.25 mM, KCl, 2.5 mM, D-glucose, 10 mM, CaCl_2_·2H_2_O, 2 mM, MgCl_2_·6H_2_O, 2 mM, adjusted to pH ∼7.4 and ∼300 mOsm). An internal cannula connected to a microsyringe (701N 10 μl, Hamilton) through silicon tube was inserted into the implanted guide cannula and muscimol (0.5-5 μl) was infused (∼1 μl/min). The internal cannula was removed carefully and the examination of eyelid movements resumed within 15 mins. Mice were imposed ≥ 30 trials before/after the infusion. In some mice (90%_CS-US_, n = 3; 10%_CS-US_, n = 3), aCSF was infused without muscimol on another day for negative control. The session was omitted from analysis when UR or ordinal behavior significantly changed after the infusion.

Muscimol diffused area and the cannula position were confirmed in individual mice. Acriflavine, a fluorescent dye, (0.5-5 μl, 0.01%, Wako), was infused in the same way as the muscimol infusion (90%_CS-US_, n = 5; 10%_CS-US_, n = 6) after pharmacological inactivation experiments. Mice were euthanized by cervical dislocation and the head plates were detached from the skulls. Whole brains were carefully removed and fixed in 4% paraformaldehyde/PBS (paraformaldehyde: Nakalai Tesque; PBS: Takara Bio) overnight and soaked for >12 hours in ∼20% sucrose for cryoprotection. Horizontal sections were prepared at ∼50 μm thickness using a cryostat (SM2000R, Leica). These sections were immediately mounted (VECTARSHIELD Mounting Medium, Funakoshi) and imaged with a confocal fluorescent microscope (FV1000, Olympus). The area of muscimol diffusion was estimated from the acriflavine fluorescence. The position of small holes in the sections was regarded as the positions of cannula.

### Modeling

Learning model for CR acquisition and extinction was constructed, and the dynamics of probability of CR occurrence was simulated under the Python programming language (Figs. 5, 6, Figs. S5, S6). Memories for CR execution (*v1*) and suppression (*v2*) were assumed to be formed independently. As for *v1*, *v1_1st_* and *v1_2nd_* with distinct sensitivity to CR extinction or suppression were assumed based on the fact that the CRs showed resistance to extinction depending on the enrichment of paired sensory experiences (see Fig. 4C), together with the previous studies suggesting two candidate synapses encoding the CR-triggering memory with distinct resistance for extinction: synapses on cerebellar PCs and those in DCN (Garcia & Mauk, 1998; Medina et al., 2001; Ohyama et al., 2006; Antonietti et al., 2016; Antonietti et al., 2017; Broersen et al., 2023). On the other hand, trials-dependent gradual suppression by the memory *v2*, was implemented by the gate control ‘*g*’, which made the spontaneous recovery possible in the model.

Taken together, the *CR_ratio_* was expressed as a function of three memories, as follows:

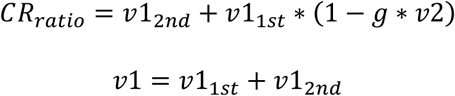

Three memories, *v1_1st_*, *v1_2nd_* and *v2*, changed trial by trial based on Rescorla-Wagner model (Rescorla & Wagner, 1972), as follows:

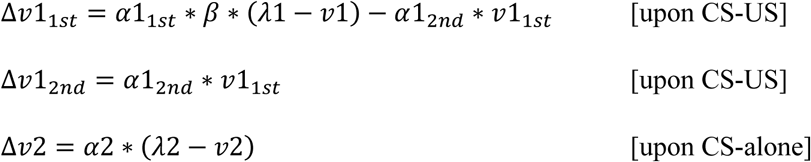

, where *λ1* and *λ2* are saturation levels for *v1* and *v2*, and set to 1 here. *α1_1st_*, *α1_2nd_*, and *α2* are rate constants for formation of *v1_1st_*, *v1_2nd_* and *v2*, respectively. *β* was assumed here to make the CS-US pair-dependent formation of *v1_1st_* sensitive to the total number of daily paired trials within a session (*N_paired_*) (Fig. S5A):

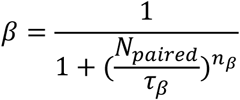

, where *τ_β_* and *n_β_* define the steepness of experience-dependent slowing of v*1_1st_* formation.

Gating of suppressing memory *v2, g*, was also implemented as follows:

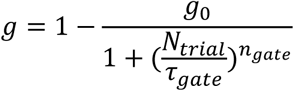

, where *N_trial_* is the total trial number in a session. *g_0_* is a factor to decide the initial gate active at the beginning of each session. *τ*_gate_ and *n_gate_* define the steepness of the trials-dependent gate opening, and the former was defined to change depending on experience as follows (Fig. S5B):

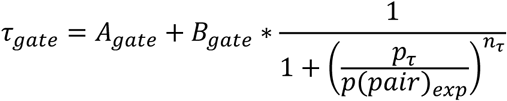

, where *A*_*gate*_ and *B*_*gate*_ are the initial value and the experience-dependent maximal increment of *τ*_gate_, respectively. *p_τ_* and *n_τ_* are constants to define the steepness of *τ_gate_* change. Expected paring ratio, *p(pair)_exp_,* was assumed to be updated trial by trial as follows (Fig. S5C):

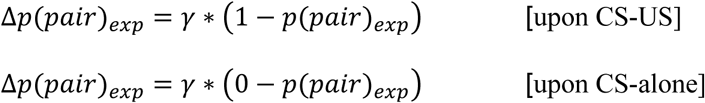

, where *γ* is the rate constant for change of *p(pair)_exp_*.

Individual memories were assumed to fade over time (*t*) as defined by the following equations:

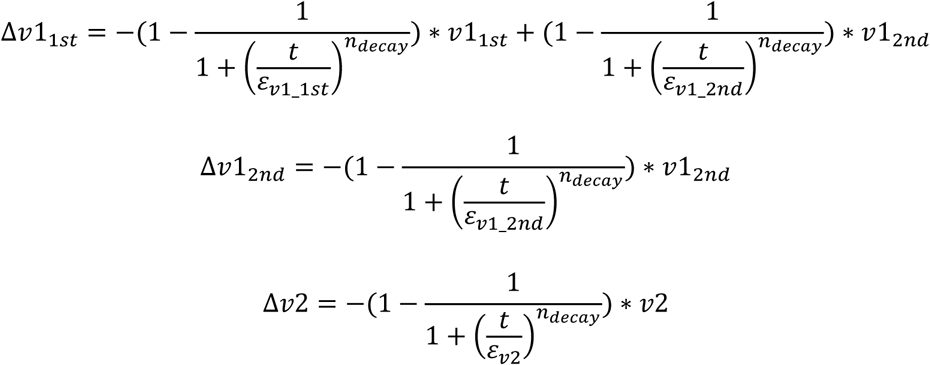

, where ε*_v1_1st_*, ε*_v1_2nd_*, *ε_v2_*, and *n_decay_* are time constants of memory decay and their steepness. Daily decay of each was calculated before each session started in simulation using 1440 min, corresponding to 24 hours, for time t. Other variables, *β*, *g*, and *p(pair)_exp_* are refreshed to initial values at the beginning of each day (see Table S1).

Simulation was performed trial by trial, giving CS-US paired or CS-alone trials as in the experiments. The 2 models assuming long- or short-lasting *v2* suppressing memory, and a model without gate control were performed by changing corresponding parameters (see Table S1).

### Statistics

Data was presented as mean ± SEM unless otherwise stated. Paired t-test and Tukey-Kramer test were performed with Microsoft Excel or Rstudio (Posit) for statistical analysis. *, p < 0.05; **, p < 0.01; ***, p < 0.001 were regarded as significant. Approximate curves were obtained with Igor Pro8 (WaveMetrics) using Chi-square method.

## Acknowledgments

We thank Drs T. Inoshita and H. Hirai for the critical reading of the manuscript and helpful comments.

## Funding

Japan Society for Promotion of Science, KAKENHI grants 22H02721 (SK)

Japan Society for Promotion of Science, KAKENHI grants 22K19360 (SK)

Japan Society for Promotion of Science, KAKENHI grants 25H02611 (SK)

Japan Society for Promotion of Science, KAKENHI grants 25K02362 (SK)

Takeda Science Foundation (SK)

JST SPRING JPMJSP2110 (RI)

## Author contributions

Conceptualization: RI, SK; Investigation: RI; Visualization: RI, SK; Supervision: SK; Writing - original draft: RI, SK; Writing – review & editing: RI, SK

## Data and materials availability

All data are available in the main text or the supplementary materials. Program codes for CR behavior simulation are also available on the website of the corresponding author (www.nb.biophys.kyoto-u.ac.jp/model).

## Supplementary materials

Figs. S1 to S7 Table S1

