## Supplementary Figures and Table for "Parallel formation of opposing memories tunes online and pre-emptive control of learned behavior in eyeblink conditioning"

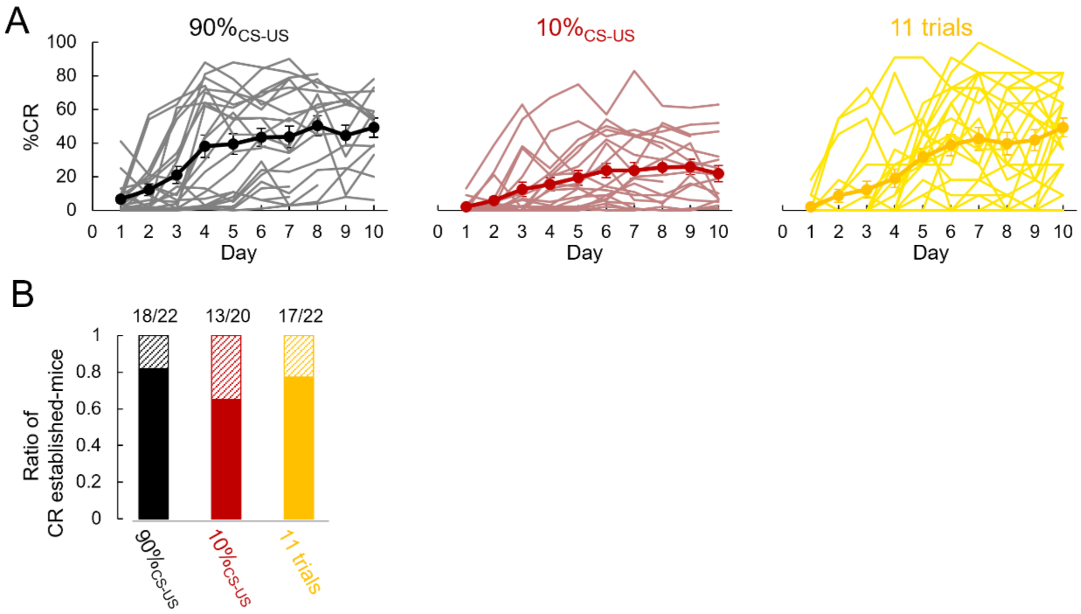


**Fig S1. CR acquisition in individual mice**

(**A**) Daily %CR changes during 10 days of training in all individual mice (individual, thin; average, bold). (**B**) Proportion of CR-established mice. n = 22, 20, 22 for the 90%$_{CS\text{-}US}$, 10%$_{CS\text{-}US}$, 11 trials, respectively.

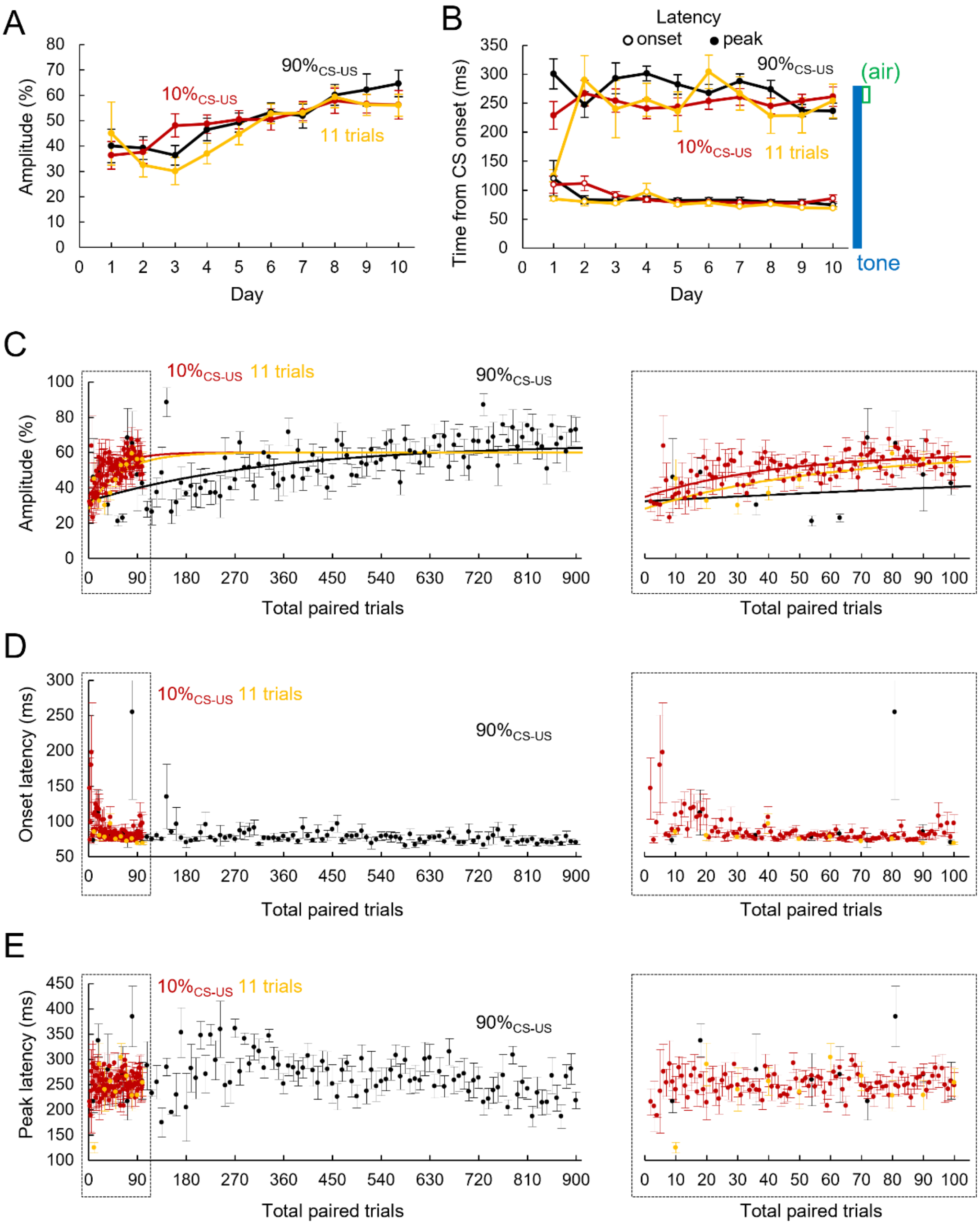


**Fig S2. Timing and amplitude changes in eyeblink responses during acquisition**

(**A**, **B**) Daily changes of averaged amplitude (A) and onset/peak latency (B) for CS-alone trials. Paired trials were also included for amplitude in the 11 trials group to compensate for small number of available traces. n = 18 (90%$_{\text{CS-US}}$), 13 (10%$_{\text{CS-US}}$), and 17 (11 trials). (**C-E**) Eyelid closure amplitude with exponential curve fits (C) and onset/peak latency (D/E) averaged from each 10 or 11 trial plotted against the total paired trials. As for fittings, initial and final amplitudes, the number of paired stimuli to reach ~63% of amplitude increment, $\tau$, and the $R^2$ for fitting were obtained as follows: initial amplitude, 32.4 (90%$_{CS-US}$), 34.8 (10%$_{CS-US}$), 28.2 (11 trials); final amplitude, 65.0 (90%$_{CS-US}$), 60.0 (10%$_{CS-US}$), 60.0 (11 trials); $\tau$, 346.0 (90%$_{CS-US}$), 43.1 (10%$_{CS-US}$), 57.3 (11 trials); $R^2$, 0.39 (90%$_{CS-US}$), 0.43 (10%$_{CS-US}$), 0.57 (11 trials).

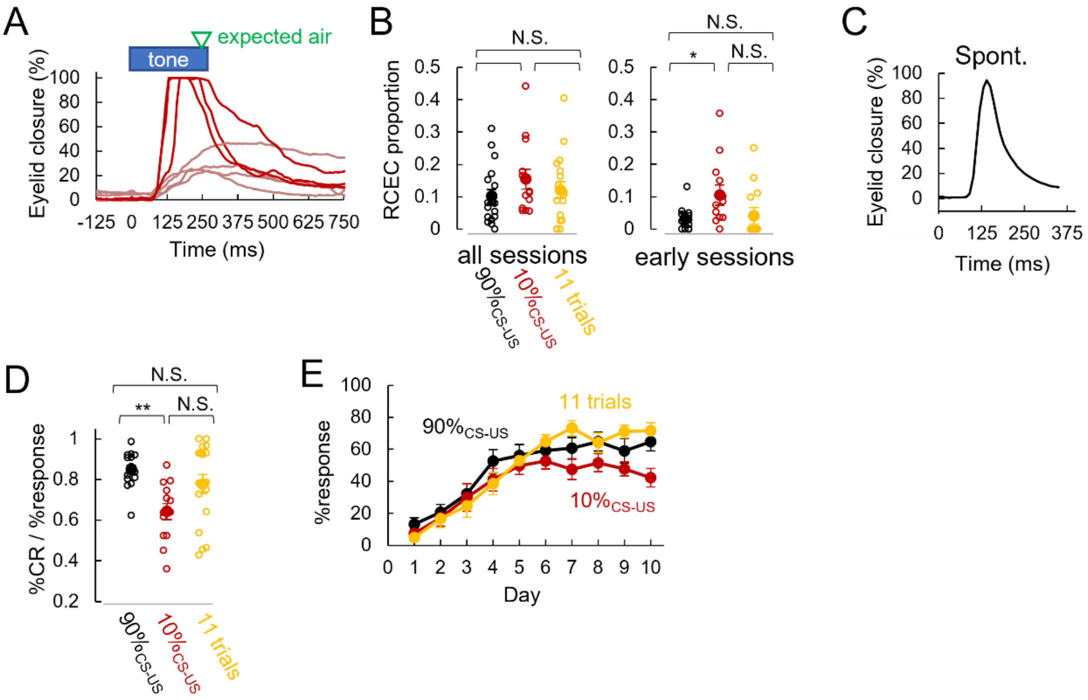


**Fig S3. Dynamics of rapid and complete eyelid closure**

(**A**) Example traces of RCEC (red) and well-timed eyeblink responses (pale red) observed from one mouse in 10%$_{\text{CS-US}}$ group. (**B**) Averaged proportion of RCEC in all/early (day1-3 or 4) sessions. Individual, open circle; average, filled circle. n = 18 (90%$_{\text{CS-US}}$), 13 (10%$_{\text{CS-US}}$), and 17 (11 trials). *, $p < 0.05$ by Tukey Kramer test. (**C**) Spontaneous eyeblink response (average of 30 traces obtained from 6 mice). (**D**) Averaged ratio of %CR to %trials with response during 10 days of training. **, $p < 0.01$ by Tukey-Kramer test. (**E**) Daily changes of %response.

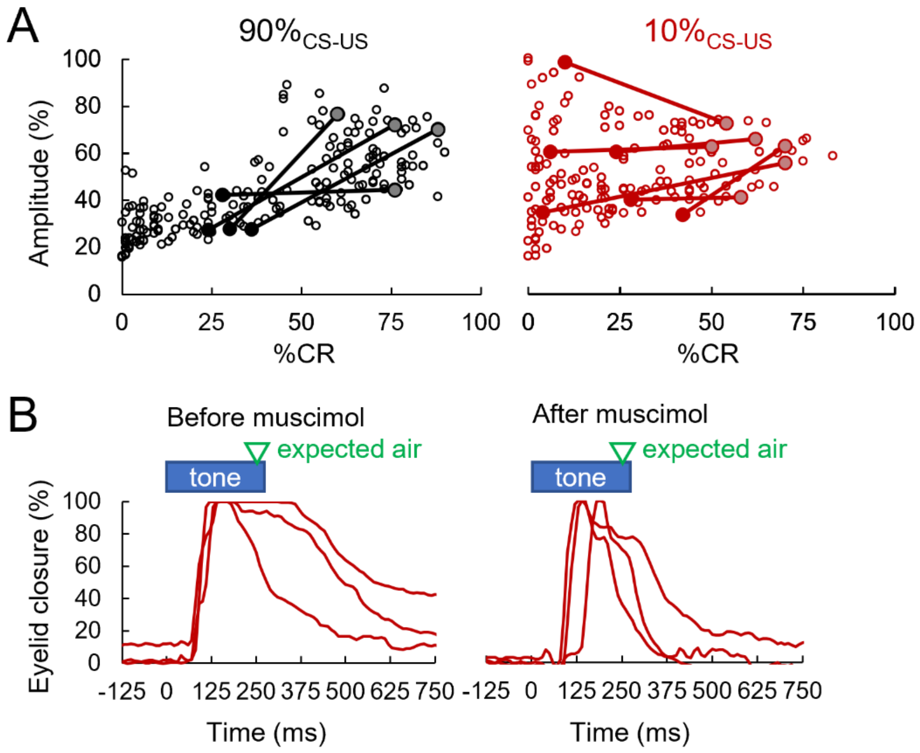


**Fig S4. Destructed relation between percentage and amplitude of CRs acquired with limited reinforcement**

(**A**) Scatter plots of averaged eyelid closure amplitude plotted against %CR for each day (open circle, $90\%_{CS\text{-}US}$, n = 24; $10\%_{CS\text{-}US}$, n = 19). Data changes before (gray or pale red)/after (black or red) muscimol infusion are highlighted by lines. n = 4 ($90\%_{CS\text{-}US}$) and n = 6 ($10\%_{CS\text{-}US}$) mice. (**B**) Representative RCEC traces observed before (left)/after (right) muscimol infusion in $10\%_{CS\text{-}US}$ group.

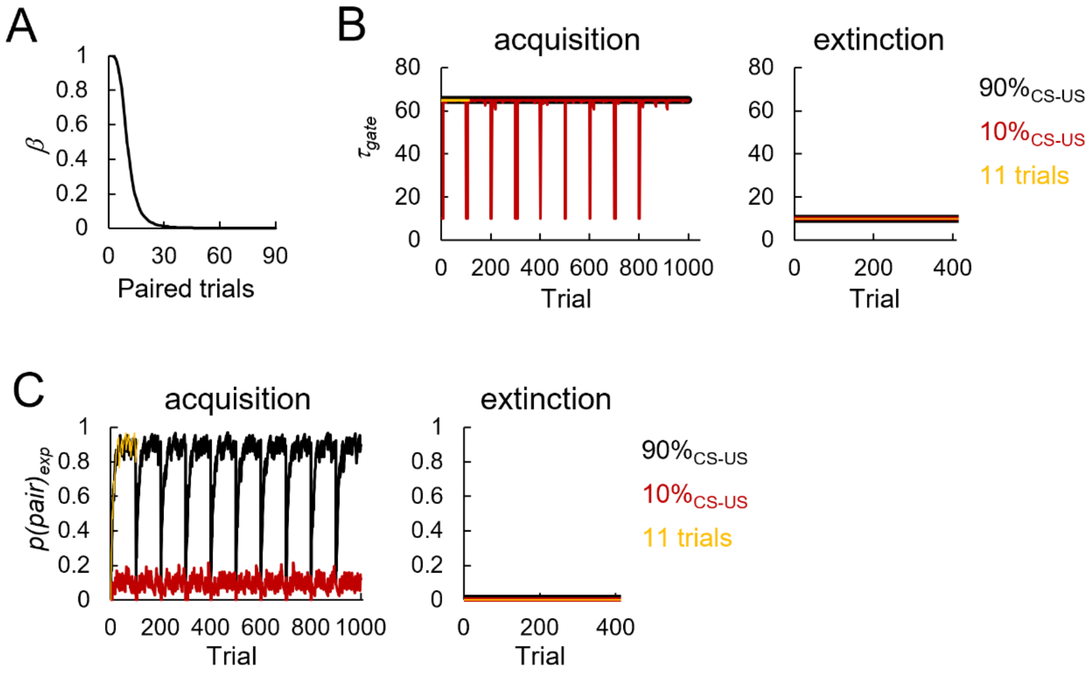


**Fig S5. Changes of variables during learning paradigm in model simulation.**

**(A)** Efficiency of $\alpha 1_{1st}$, $\beta$, plotted against the number of paired trials. **(B, C)** $\tau_{gate}$ (B) and $p(pair)_{exp}$ (C) were plotted against total trial number during acquisition (left) and extinction (right) paradigms.

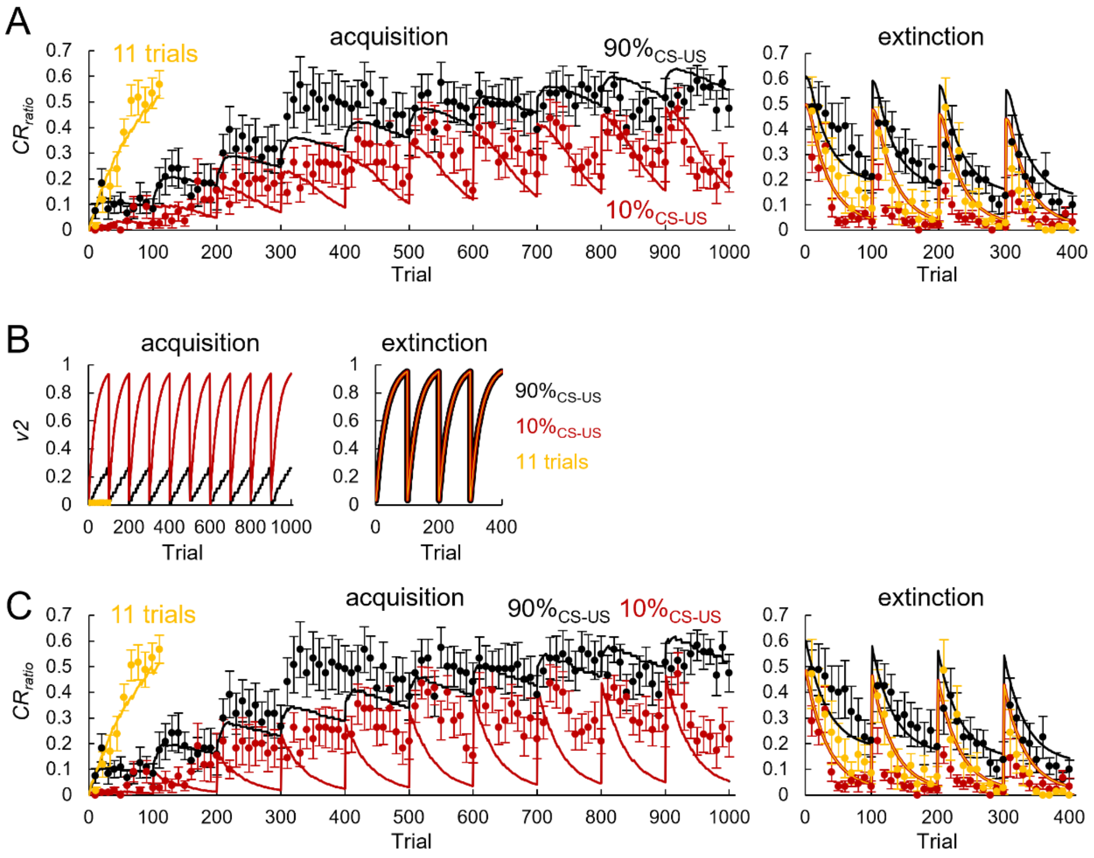


**Fig S6. CR acquisition and extinction simulated in a model assuming short-lasting v2 memory**

(**A**) Simulated $CR_{ratio}$ changes in a model with short-lasting suppressive memory, *v2,* during acquisition (left) and extinction (right) paradigms. For comparison, %CR averaged from each 10 or 11 trial recorded in experiments are also presented (dots). n = 12 (90%$_{\text{CS-US}}$), 11 (10%$_{\text{CS-US}}$) and 16 (11 trials). **(B)** Simulated changes of simulated suppressing memory, *v2*, during acquisition (left) and extinction (right) plotted against total trial number. **(C)** Simulation results obtained when the model was assumed to lack 'gate' (i.e. g was set to be 1).

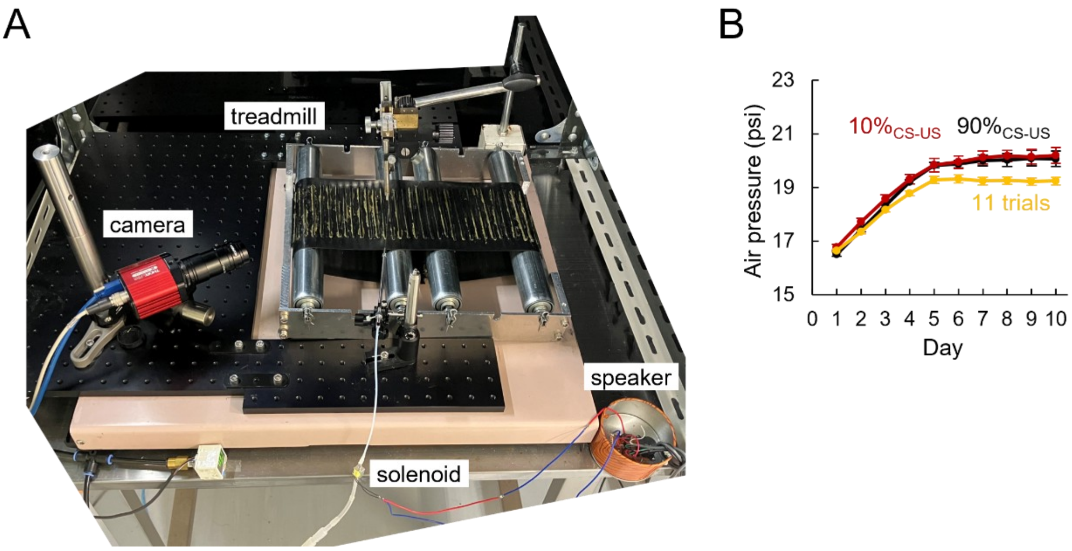


**Fig S7. Apparatus for eyeblink conditioning.**

**(A)** A photograph of the self-developed setup for eyeblink conditioning. **(B)** Averaged air intensity during 10 days of trainings given to the CR established mice. n = 18, 13, 17 mice for the $90\%_{CS\text{-}US}$, $10\%_{CS\text{-}US}$, 11 trials, respectively

| category | parameter | model1 (long) | model2 (short) | Model2-2 (no gate) |
|---|---|---|---|---|
| learning ratio | $\alpha 1_{1st}$ | 0.0105 | 0.0105 | 0.0105 |
| | $\alpha 1_{2nd}$ | 0.001 | 0.001 | 0.001 |
| | $\alpha 2$ | 0.009 | 0.03 | 0.03 |
| efficiency of $\alpha 1_{1st}$ ($\beta$) | $\tau_\beta$ | 10 | 10 | 10 |
| | $n_\beta$ | 4 | 4 | 4 |
| gate | $g_0$ | 0.8 | 0.8 | - |
| | $n_{gate}$ | 2.1 | 2.1 | - |
| | $A_{gate}$ | 10 | 10 | - |
| | $B_{gate}$ | 55 | 55 | - |
| | $p_\tau$ | 0.01 | 0.01 | - |
| | $n_\tau$ | 3 | 3 | - |
| | $\gamma$ | 0.1 | 0.1 | - |
| memory decay | $\varepsilon_{v1\_1st}$ | 7000 | 7000 | 7000 |
| | $\epsilon_{v1\_2nd}$ | 4000 | 4000 | 4000 |
| | $\varepsilon_{v2}$ | 8000 | 100 | 100 |
| | $n_{decay}$ | 2 | 2 | 2 |

| | Initial values |
|---|---|
| $v1_{1st}$ | 0 |
| $v1_{2nd}$ | 0 |
| $v2$ | 0 |
| $\beta$ | 1 |
| $g$ | 0.2 |
| $p(pair)_{exp}$ | 0 |

**Table S1. Parameters and initial values for model simulation.**
